# Paraptosome: A Novel Pathological Feature in Paraptotic Cell Death

**DOI:** 10.1101/2024.08.07.606501

**Authors:** Xiang Cui, Hongda Zheng, Haoming Li, Fang Zhang, Liao Yang, Jiayu Ni, Dengfeng Wang, Huali Zhang, Pan Tang, Ru Li, Qi Zhang, Min Cui

## Abstract

Paraptosis is a novel form of programmed cell death characterized by distinct morphological features such as swelling of the endoplasmic reticulum and mitochondria, and cytoplasmic vacuolation. Unlike apoptosis, paraptosis does not involve the activation of caspases or DNA fragmentation. These unique features make paraptosis an intriguing target for cancer therapy, particularly against apoptosis-resistant cells. Here, we report a novel morphological feature of paraptosis: the formation of high-density spherical structure, which we tentatively term “paraptosome.” We found that these putative paraptosomes originate from the Golgi apparatus, appearing as high-density formations under light microscopy and colocalizing with the trans-Golgi marker β4GALT1-RFP. Time-lapse confocal microscopy and immunostaining demonstrated that putative paraptosomes form due to Golgi stress or disintegration, leading to severe disruption of Golgi function. Furthermore, we show that paraptosis inducers such as glabridin, morusin, and honokiol can cause significant alterations in the endoplasmic reticulum, mitochondria, autophagosomes, and lysosomes in U251MG glioblastoma cells; however, the formation of putative paraptosomes is not induced by isolated stress inducers. Collectively, these findings suggest that the putative paraptosome may be a novel characteristic structure of paraptosis. The discovery of paraptosomes provides a unique marker for defining paraptotic cell death and offers new insights into the characteristic pathological phenomena associated with multiple organelle dysfunction. This finding broadens the scope of cell biology research by introducing a new structural paradigm linked to paraptosis and may have implications for developing targeted therapies against apoptosis-resistant cancers.

## Introduction

Cell death is a complex biological process crucial for tissue homeostasis, development, and disease defense. Alongside well-known mechanisms like apoptosis and necrosis, various atypical cell death pathways have been elucidated in recent years [1]. Among these, paraptosis has emerged as a unique form of programmed cell death, garnering increasing attention for its potential applications in cancer therapy [2–3].

Paraptosis distinguishes itself as a unique form of programmed cell death, characterized by extensive cytoplasmic vacuolation believed to originate from the swelling of the endoplasmic reticulum (ER) and mitochondria [3–4]. Unlike apoptosis, which involves caspase activation and DNA fragmentation, paraptosis is caspase-independent and does not involve nuclear breakdown. These distinct features make paraptosis particularly intriguing for targeting cancer cells that have developed resistance to conventional apoptosis-inducing therapies.

Recent studies have demonstrated that various natural compounds, including glabridin, morusin, honokiol, and curcumin, can induce paraptosis in different tumor cells [5–8]. These compounds share common effects such as inducing cytoplasmic vacuolation, increasing ER stress responses, and inhibiting cell proliferation. At the molecular level, it typically involves the activation of MAP kinases, particularly MEK-2/ERK2, which are crucial for initiating the paraptotic process [9–12]. The disruption of internal ion homeostasis, especially an imbalance in intracellular calcium levels, is often observed and can lead to mitochondrial swelling and dysfunction [13–16]. This, in turn, contributes to the energy depletion commonly seen in paraptotic cells [4]. A key characteristic of paraptosis is the accumulation of proteins and increased ER stress [17–18]. This is often due to impaired protein degradation, such as proteasome inhibition, or interference with protein folding [19–20]. Notably, protein synthesis inhibitors can block paraptosis, suggesting a critical role for new protein synthesis in this cell death pathway [7, 9]. Another common feature is the increased generation of reactive oxygen species (ROS), which is potentially linked to mitochondrial dysfunction [21–23]. Recent studies have also implicated the involvement of certain developmental pathways, such as Wnt signaling, in the regulation of paraptosis [24–25]. Despite these insights, the precise molecular mechanisms driving these events and the interactions among involved organelles remain poorly understood, highlighting the need for further investigation into this unique form of cell death.

In our preliminary studies using paraptosis inducers, we observed the formation of novel high-density spherical structure in the cytoplasm of treated cells. We hypothesize that the structure, which we tentatively term “paraptosome,” may represent a previously unrecognized feature of paraptosis and may play a unique and intriguing role in this process.

This study aims to characterize the morphology, composition, and dynamics of paraptosomes in relation to known cellular organelles during paraptosis. We seek to elucidate their origin, with a particular focus on their potential relationship to the Golgi apparatus, which has been understudied in the context of paraptosis. By investigating the effects of paraptosis inducers on cellular organelle function, protein trafficking, and stress responses, we aim to evaluate the potential of paraptosomes as a distinctive morphological marker for paraptosis and examine their functional implications in cellular stress responses and cell death mechanisms.

To achieve these objectives, we employ a combination of advanced cell biology techniques, including specific fluorescent organelle trackers, time-lapse confocal microscopy, and transmission electron microscopy. These methods allow for precise delineation of organelle dynamics and interactions during the paraptotic process. Additionally, we utilize biochemical assays and gene expression analysis to elucidate the functional changes associated with these structures.

By elucidating the nature and significance of these novel structures, we expect to provide new insights into the mechanisms of paraptosis. This research may not only enhance our understanding of this unique cell death pathway but also lay the groundwork for developing innovative therapeutic strategies against drug-resistant cancers.

## Results

### Identification of High-Density Spherical Structures in Paraptosis Induced by Natural Compounds

While extensive cytoplasmic vacuolation is a well-established hallmark of paraptosis, our investigation of glabridin-induced paraptosis in breast cancer cells revealed novel high-density spherical structures alongside typical low-density vacuoles [5]. To further explore this phenomenon, we extended our study to include several natural compounds known to induce paraptosis: glabridin, morusin, honokiol, and curcumin [5–8].

We first assessed the cytotoxicity of these compounds in U251MG glioblastoma cells. CCK-8 assays revealed a dose-dependent decrease in cell viability after 24-hour treatment with concentrations ranging from 20 μM to 100 μM (Figure 1A). Flow cytometry with PI staining showed a significant increase in PI-positive cells in treated groups, indicating compromised cell membrane integrity increased cell death (Figure 1B). Notably, western blot analysis revealed no significant increase in the cleavage of caspase-3 and PARP, two key markers of apoptosis, in cells treated with glabridin, honokiol, and morusin (Figure 1C). However, curcumin-treated cells exhibited a significant increase in cleaved PARP, suggests curcumin may involve a distinct mechanism in U251MG cells.

**Figure 1:**
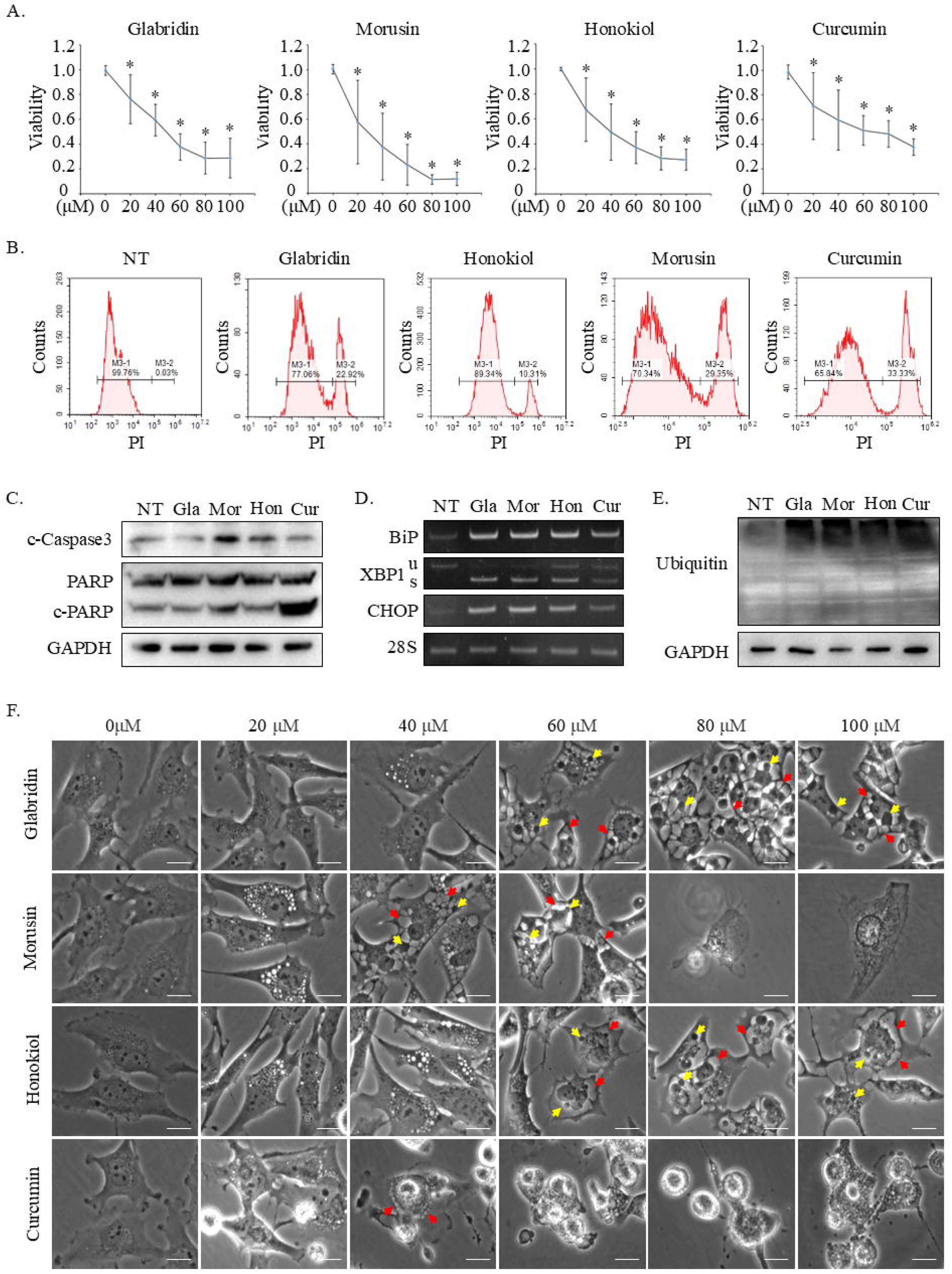
Characterization of Paraptosis Induced by Natural Compounds in U251MG Glioblastoma Cells. (A) CCK-8 assay showing dose-dependent effects of glabridin, morusin, honokiol, and curcumin (20-100 μM, 24h) on U251MG cell viability. Mean ± SD, n=3. *P<0.05 vs. control. (B-E) U251MG cells treated with glabridin (60 μM), morusin (40 μM), honokiol (60 μM), or curcumin (40 μM) for 24h. (B) Cell death percentage assessed by PI staining and flow cytometry. (C) Western blot analysis of apoptosis markers (cleaved caspase-3 and cleaved PARP) showing no significant increase. (D) RT-PCR analysis of ER stress markers (BiP, spliced XBP1, and CHOP) indicating differential upregulation. 28S was used as a loading control. (E) Western blot showing accumulation of ubiquitinated proteins. GAPDH served as a loading control. (F) Light microscopy images illustrating morphological changes in U251MG cells treated for 24h with 20-100 μM glabridin, morusin, honokiol, or curcumin. Red arrows indicate low-density vacuoles, while yellow arrows indicate high-density spherical structures (paraptosomes). Scale bar: 20 μm. NT: Non-Treatment, Gla: Glabridin, Mor: Morusin, Hon: Honokiol, Cur: Curcumin, PI: Propidium Iodide

Additionally, we assessed two other known hallmarks of paraptosis: ER stress and proteasome inhibition. RT-PCR analysis showed differential upregulation of ER stress markers (BiP, spliced XBP1, and CHOP) [26] in response to the tested compounds. Glabridin, morusin, and honokiol treatment resulted in pronounced increases in these markers, whereas curcumin treatment led to a comparatively milder upregulation (Figure 1D). Furthermore, Western blot analysis revealed a substantial accumulation of ubiquitinated proteins induced by all four compounds (Figure 1E).

Under light microscopy, we observed distinct morphological changes in U251MG cells treated with glabridin, morusin, and honokiol (Figure 1F). At concentrations of 60 μM for glabridin, 40 μM for morusin, and 60 μM for honokiol, cells exhibited the formation of typical low-density vacuoles (red arrows) and high-density spherical structures (yellow arrows). Furthermore, the size of these high-density structures increased with higher drug concentrations. In contrast to the other compounds, curcumin showed a distinct response pattern. At lower concentrations, curcumin treatment resulted in the formation of small cytoplasmic vacuoles (Figure 1F and Supplementary Figure S1). However, at concentrations above 40 μM, curcumin caused significant cell shrinkage and reduced cell numbers, preventing the observation of enlarged vacuoles or high-density spherical structures seen with the other compounds. This distinct behavior suggests curcumin may operate via a different mechanism in U251MG cells (Figure 1F).

These findings suggest that the high-density spherical structure, which we tentatively term “paraptosome” for convenience in subsequent studies, may be a characteristic morphological change associated with paraptosis induced by certain compounds.

### ER and Mitochondrial Alterations during Paraptosis

Previous research has shown that cytoplasmic vacuoles in paraptosis originate from the ER or mitochondria [3, 5]. To elucidate whether putative paraptosomes also originate from these organelles, we employed specific fluorescent trackers: KDEL-GFP for the ER and COX8-RFP for mitochondria. U251MG cells stably expressing KDEL-GFP, and COX8-RFP were treated with glabridin (60 μM), morusin (40 μM), honokiol (60 μM), and curcumin (40 μM) for 24 hours. Under normal conditions, KDEL-GFP exhibited a reticular distribution characteristic of the ER network throughout the cytoplasm (Figure 2A). However, treatment with glabridin, morusin, and honokiol caused the green-fluorescent signals to aggregate in areas of cytoplasmic vacuolation, suggesting a relocalization of the ER (red arrows). COX8-RFP fluorescence revealed disrupted mitochondrial networks in treated cells, with mitochondria appearing as dispersed punctate or clumped structures (Figure 2B). Notably, these clumped regions did not overlap with putative paraptosomes, suggesting putative paraptosomes are not derived from fragmented mitochondria.

**Figure 2:**
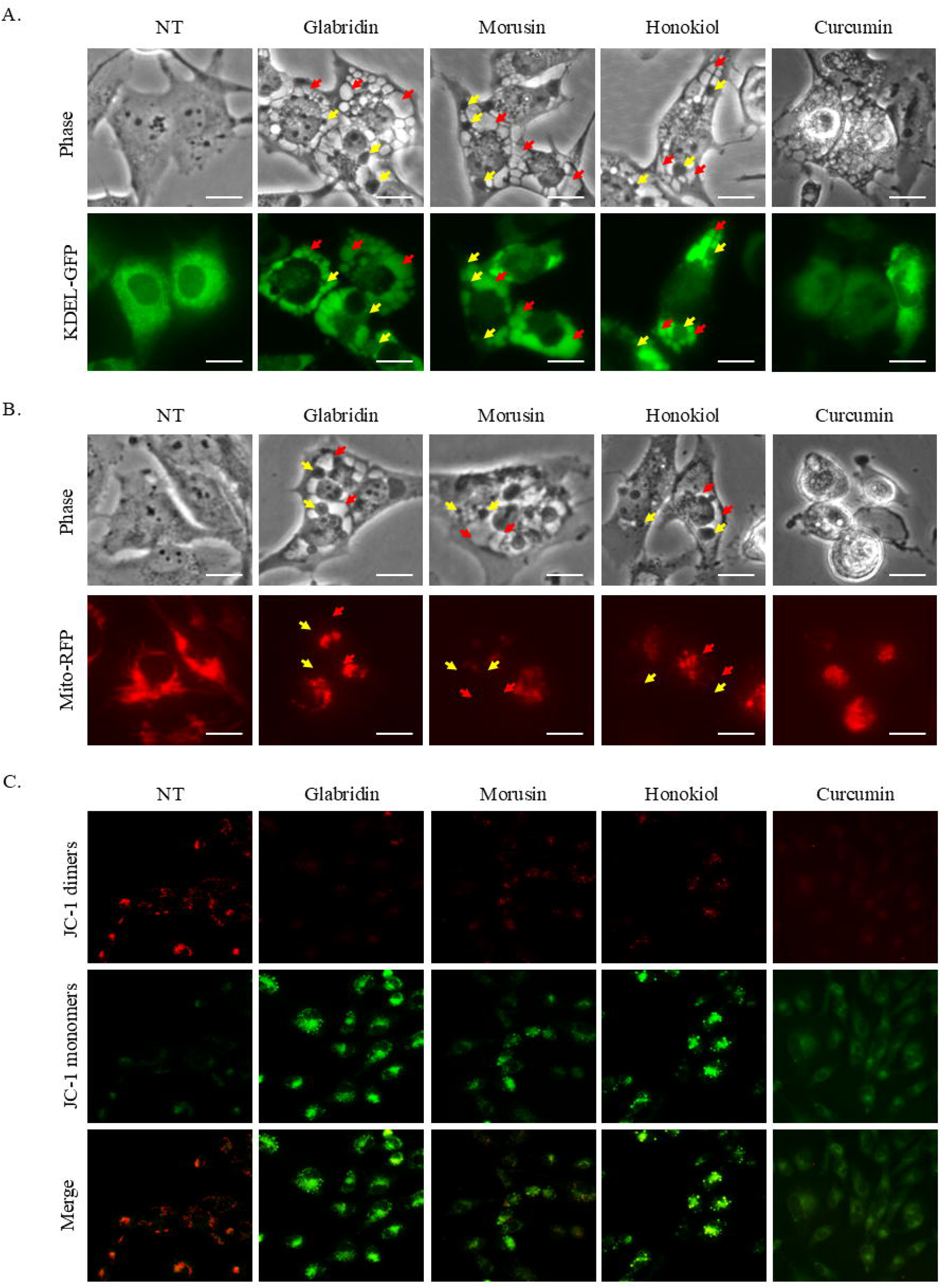
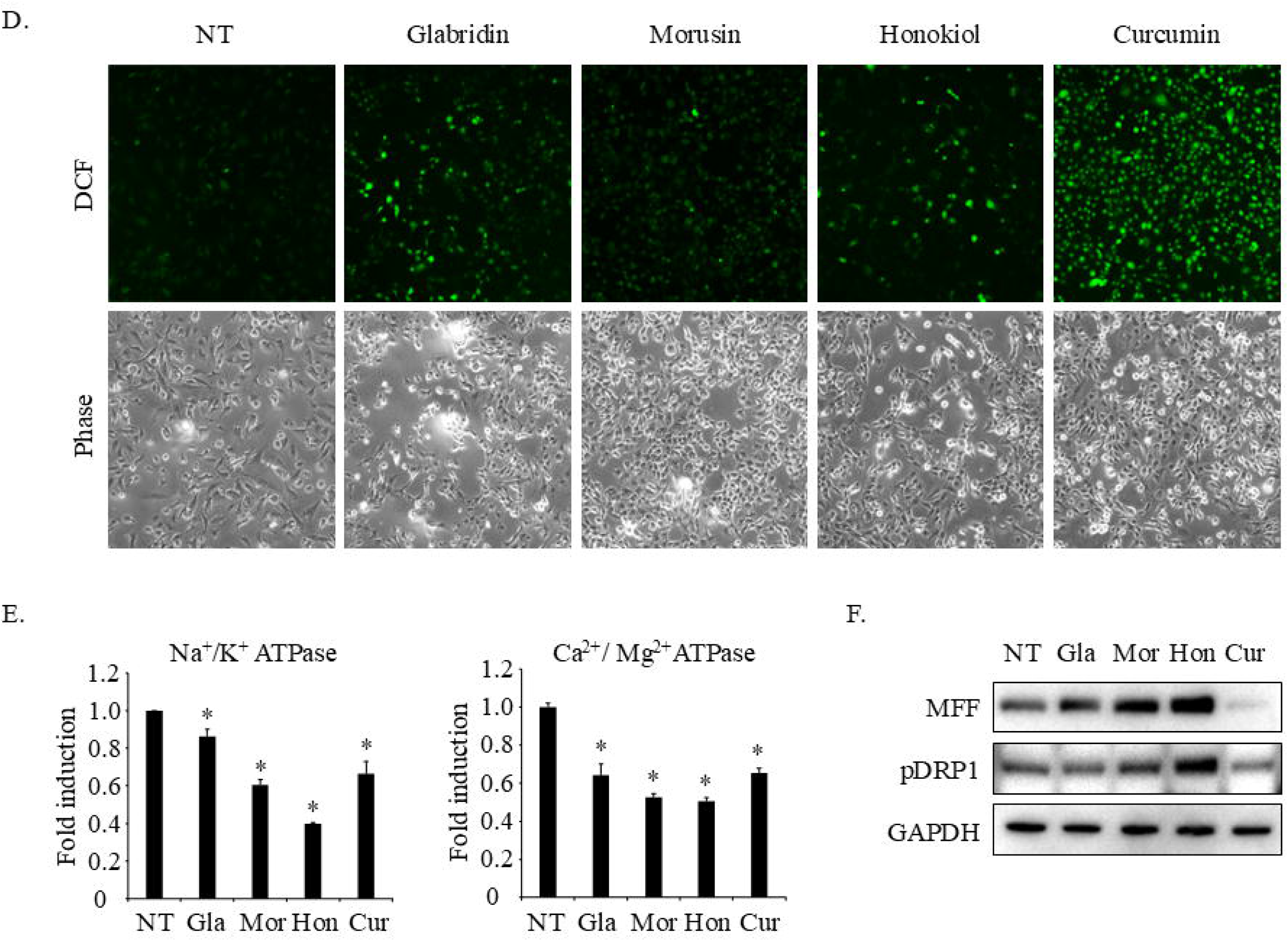
Paraptosomes Arise Independently of ER and Mitochondrial Alterations. (A) U251MG cells stably expressing KDEL-GFP treated with glabridin (60 μM), morusin (40 μM), honokiol (60 μM), curcumin (40 μM) for 24 hours. ER signals aggregate in areas of cytoplasmic vacuolization (red arrows) but did not overlap with high-density paraptosomes (yellow arrows). Scale bar: 20 μm. (B) Fluorescent microscopy images of U251MG cells stably expressing COX8-RFP treated as in (A). Disrupted mitochondrial networks appear as dispersed punctate or clumped structures, that did not overlap with the putative paraptosomes (yellow arrows). Scale bar: 20 μm. (C-F) U251MG cells treated with glabridin (60 μM), morusin (40 μM), honokiol (60 μM), or curcumin (40 μM) for 6h. (C) JC-1 staining indicating loss of mitochondrial membrane potential with increased green fluorescence. (D) ROS levels measured by DCFH-DA fluorescence showing slight increases with glabridin, morusin, and honokiol, and a substantial increase with curcumin. (E) Na^+^/K^+^ and Ca^2+^/Mg^2+^ ATPase assays showing significant decrease in all treated cells. Mean ± SD, n=3. *P<0.05 vs. control. (F) Western blot analysis showing upregulation of MFF and Phospho-DRP1 (pDRP1) in cells treated with glabridin, morusin, and honokiol, but not curcumin. GAPDH was used as a loading control.

Further analysis of mitochondrial function revealed significant changes. JC-1 staining showed a loss of mitochondrial membrane potential in cells treated with glabridin, morusin, and honokiol, as evidenced by increased green fluorescence (Figure 2C). ROS levels slightly increased with glabridin, morusin, and honokiol treatments, while curcumin caused the most substantial increase (Figure 2D). All four compounds significantly decreased Na^+^/K^+^ and Ca^2+^/Mg^2+^ ATPase activities (Figure 2E), suggesting a broad impact on cellular energy metabolism and ion homeostasis. Western blot analysis demonstrated that glabridin, morusin, and honokiol treatments led to the upregulation of mitochondrial dynamics proteins MFF and Phospho-DRP1 (Figure 2F). In contrast, curcumin did not induce similar changes, indicating a distinct mode of action compared to the other compounds.

These findings elucidate a distinct reorganization of ER and mitochondrial morphology and function in response to paraptotic inducers, yet putative paraptosomes appear to arise independently of these organelles.

### Lysosomal and Autophagic Dynamics in Paraptosis

To elucidate the origin and nature of putative paraptosomes, we investigated their relationship with lysosomes and autophagosomes. U251MG cells stably expressing the lysosomal marker Lamp1-RFP or the autophagy marker RFP-LC3B were treated with glabridin (60 μM), morusin (40 μM), honokiol (60 μM), and curcumin (40 μM) for 24 hours. Following treatment, the Lamp1-RFP signal distribution changed significantly from a dispersed circular pattern to irregular, clustered aggregates (Figure 3A). Similarly, RFP-LC3B fluorescence patterns were altered but did not coincide with cytoplasmic vacuoles or putative paraptosomes (yellow arrows, Figure 3B). Immunostaining for p62, a key autophagy regulator that binds to LC3 and ubiquitinated substrates, showed a marked redistribution from a diffuse cytoplasmic pattern in control cells to distinct punctate or aggregated structures in treated cells (Figure 3C). It is important to note that during immunostaining, a significant proportion of putative paraptosomes disappeared after paraformaldehyde (PFA) fixation, which may affect the detection of these structures. This observation suggests that putative paraptosomes may be sensitive to standard fixation protocols, highlighting the need for alternative methods to preserve these structures for immunofluorescence studies.

**Figure 3:**
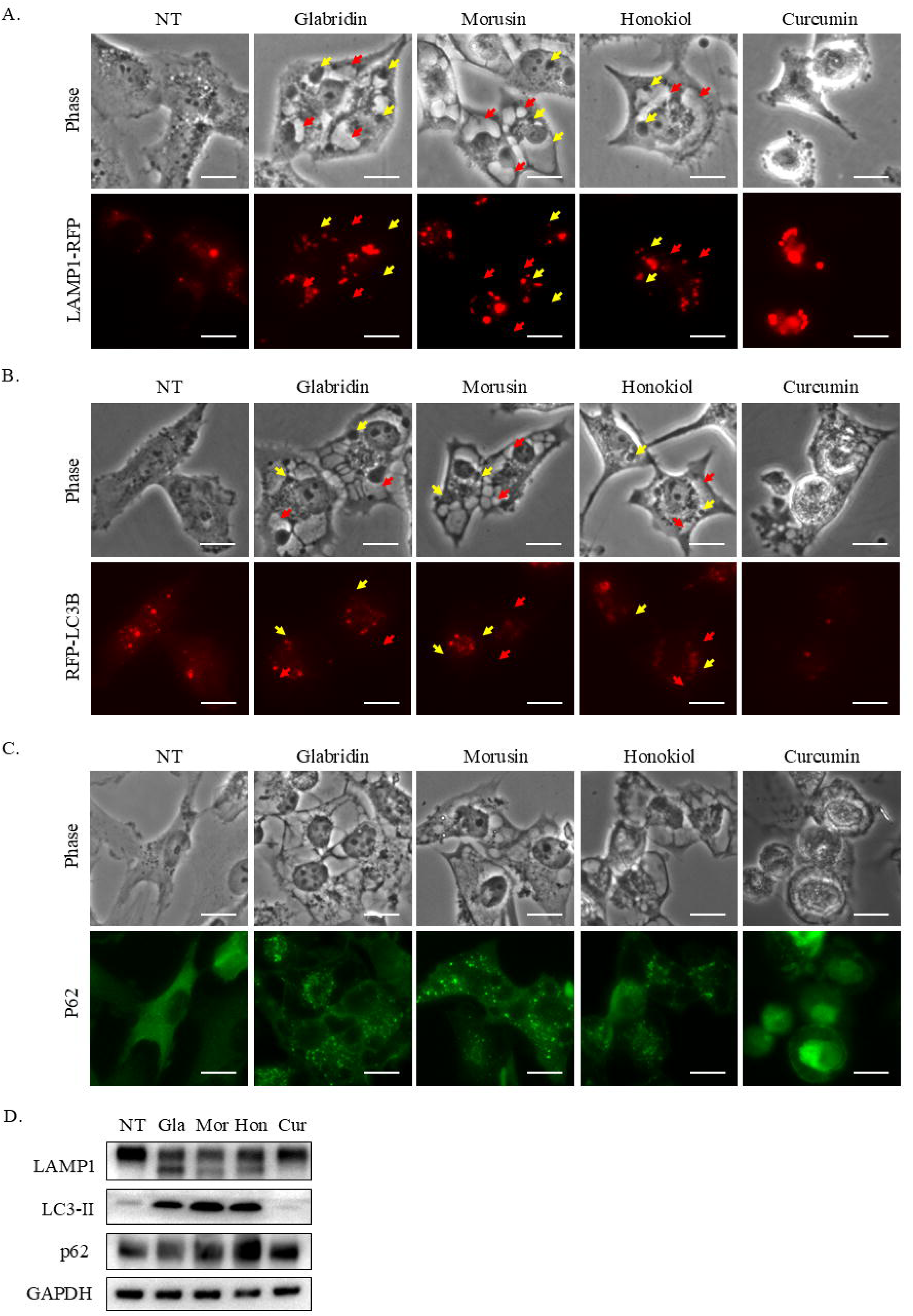
Lysosomal and Autophagic Alterations During Paraptosis. (A) U251MG cells stably expressing Lamp1-RFP treated with glabridin (60 μM), morusin (40 μM), honokiol (60 μM), and curcumin (40 μM) for 24 hours. Lamp1-RFP signal distribution changed from a dispersed circular pattern to irregular, clustered aggregates. Red arrows indicate cytoplasmic vacuoles, and yellow arrows indicate paraptosomes. Scale bar: 20 μm. (B) U251MG cells stably expressing RFP-LC3B treated as in (A). RFP-LC3B fluorescence patterns were altered but did not overlap with cytoplasmic vacuoles or paraptosomes. Scale bar: 20 μm. (C) Immunostaining for p62 in U251MG cells treated as in (A) showed a marked redistribution from a diffuse cytoplasmic pattern to distinct punctate or aggregated structures. Scale bar: 20 μm. (D) Western blot analysis showing a downward shift in LAMP1 protein molecular weight and increased LC3-II expression in cells treated with glabridin, morusin, and honokiol, but not curcumin. p62 expression increased only in honokiol-treated cells. GAPDH was used as a loading control.

Western blot analysis revealed a noticeable downward shift in LAMP1 protein molecular weight following treatment with glabridin, morusin, and honokiol, suggesting alterations in its post-translational modification (Figure 3D). LC3-II expression was significantly increased in cells treated with these compounds, while p62 expression increased only in honokiol-treated cells. Notably, curcumin treatment did not induce similar changes in LC3-II or p62 expression. These results suggest that putative paraptosomes are likely not derived from autophagosomes or lysosomes. However, the process of paraptosis appears to significantly disrupt lysosomal functions and autophagic processes to some extent, highlighting the complex nature of cellular responses during paraptosis.

### Golgi Apparatus Disintegration Leads to Putative Paraptosome Formation

To monitor structural changes in the Golgi apparatus during paraptosis, we utilized U251MG cells stably expressing the Golgi-specific fluorescent marker β1,4-galactosyltransferase (B4GALT1)-RFP, which is primarily localized in the trans-Golgi cisternae [27]. Under normal conditions, B4GALT1-RFP signals appeared as clustered punctate structures near the nucleus (Figure 4A). Following treatment with glabridin (60 μM), morusin (40 μM), and honokiol (60 μM) for 24 hours, we observed the formation of larger circular structures that corresponded to the putative paraptosomes observed under phase contrast microscopy. Interestingly, time-lapse confocal microscopy revealed that one hour after glabridin treatment, small circular structures began to form around the original Golgi apparatus, progressively expanding and fusing over time (Figure 4B and Supplementary Video). To gain further insights into the ultrastructure of putative paraptosomes, we performed electron microscopy analysis. The results showed that these structures are approximately spherical, composed of a dense network of intertwined membranous structures, separated from the surrounding cytoplasm by clearly defined edges (Figure 4C). Unlike typical cellular vesicles or organelles, these structures exhibited a unique morphology, characterized by their high electron density and complex internal organization.

**Figure 4:**
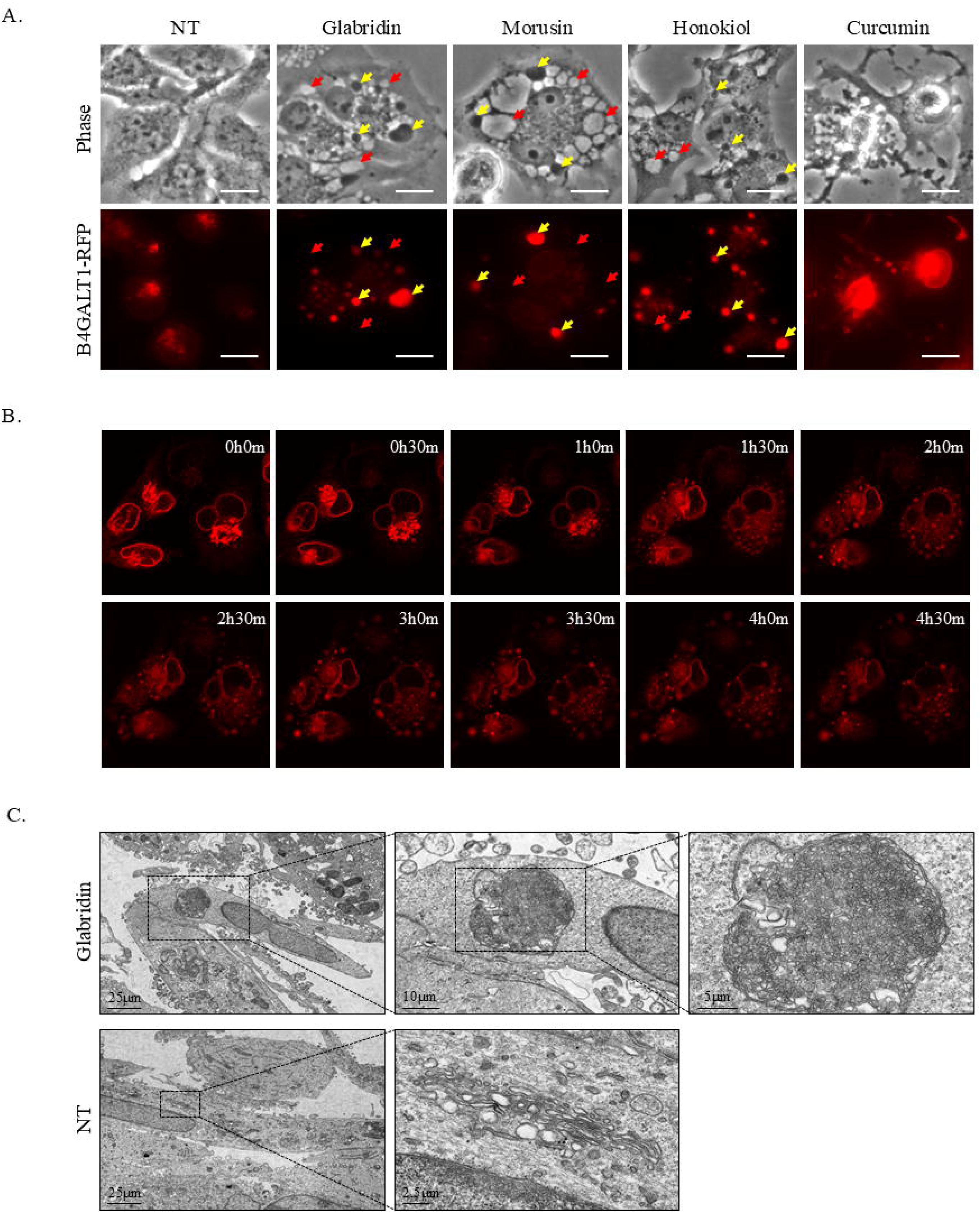
Structural Changes in the Golgi Apparatus and Paraptosome Formation During Paraptosis. (A) Phase contrast and fluorescence microscopy images of U251MG cells stably expressing B4GALT1-RFP (Golgi marker) treated with glabridin (60 μM), morusin (40 μM), honokiol (60 μM), and curcumin (40 μM) for 24 hours. Red arrows indicate low-density vacuoles, and yellow arrows indicate high-density spherical structures (paraptosomes). Scale bar: 20 μm. (B) Time-lapse confocal microscopy images of U251MG cells stably expressing B4GALT1-RFP showing dynamic changes in the Golgi apparatus after treatment with 60 μM glabridin. Images were captured at the indicated time points, demonstrating the progressive formation and expansion of paraptosomes. (C) Electron microscopy images of U251MG cells. Top row: Cells treated with glabridin (60 μM) for 24 hours. The right panels show magnified views of the dashed boxed areas in the left images, highlighting paraptosome with a dense network of intertwined membranes. Bottom row: Non-treatment (NT) control cells. The dashed boxed area in the left image highlights the normal Golgi apparatus, with the right panel showing a magnified view. Scale bars: 25 μm, 10 μm, 5 μm, 2.5μm.

These findings suggest that putative paraptosomes are likely secondary structures formed during the disintegration of the Golgi apparatus in paraptosis, highlighting the complex cellular alterations associated with this form of cell death. The formation of putative paraptosomes from Golgi-derived materials represents a previously undescribed feature of paraptosis and may provide new insights into the mechanisms underlying this cell death pathway.

### Alterations in Protein Trafficking and Golgi Function During Paraptosis

During paraptosis, significant morphological changes in the Golgi apparatus were observed, prompting an investigation into the functional impact of these alterations. Immunostaining with the cis-Golgi marker GM130 and the cis-/medial Golgi marker Giantin revealed that cells treated with paraptosis inducers exhibited complete disintegration of the typical perinuclear Golgi structure, indicating severe disruption of Golgi organization (Figure 5A and 5B). However, western blot results for GM130 and Syntaxin 16 (a trans-Golgi marker) showed no significant changes in the expression levels (Figure 5C), suggesting that the observed structural changes are due to reorganization rather than degradation of these proteins. To assess the functional impact of these structural changes, we measured the extracellular secretion of VEGFA, a protein that requires processing and secretion through the Golgi apparatus [28]. ELISA assays demonstrated a significant reduction in VEGFA secretion following treatment with paraptosis inducers (Figure 5D). Furthermore, we examined the mRNA expression levels of Golgi stress markers ARF4, ATF3, and CREB3 [29–30]. RT-PCR results showed that glabridin, morusin, and honokiol significantly induced the expression of these genes, whereas curcumin had a less pronounced effect (Figure 5E), indicating differential activation of Golgi stress responses.

**Figure 5:**
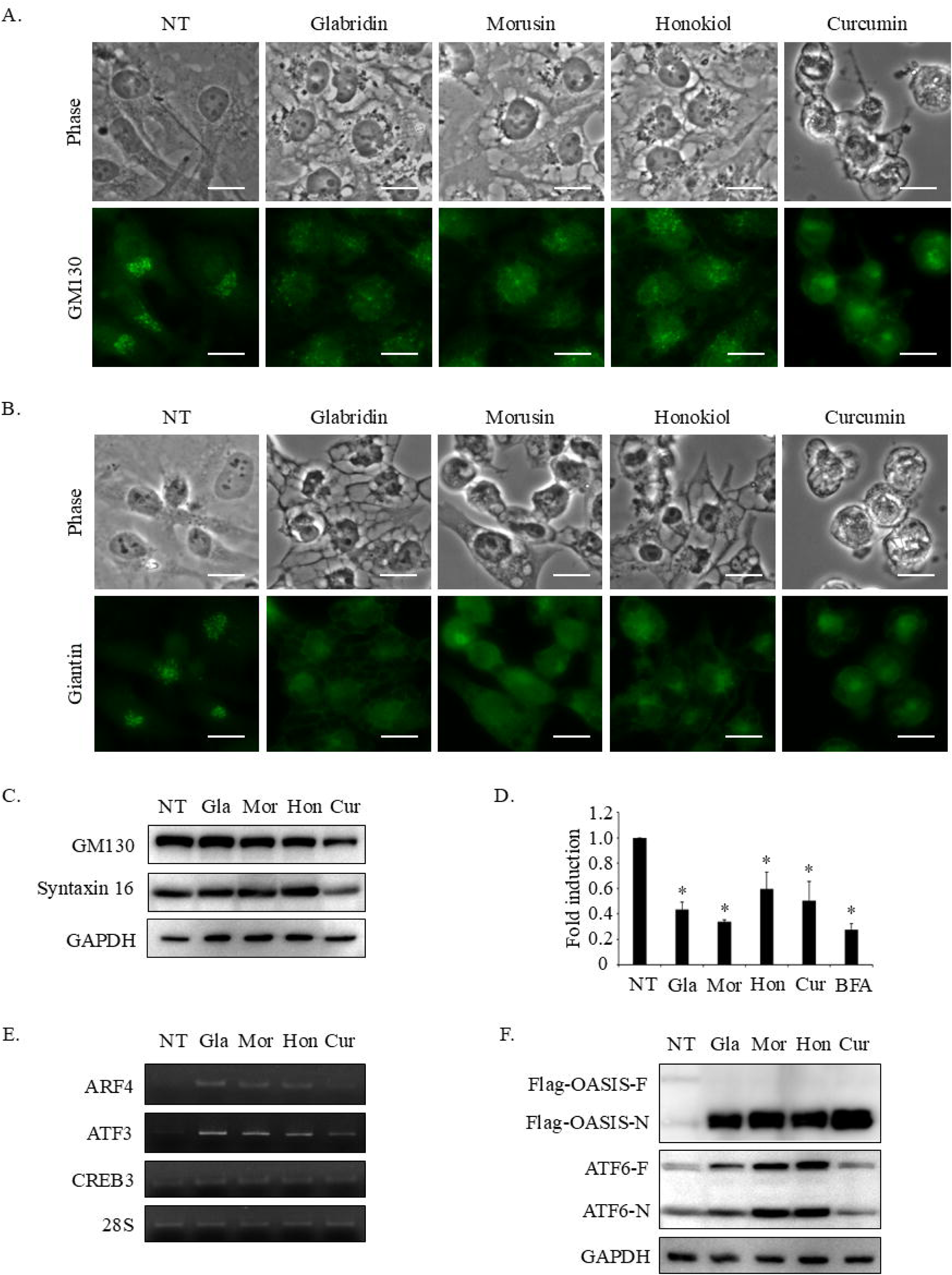
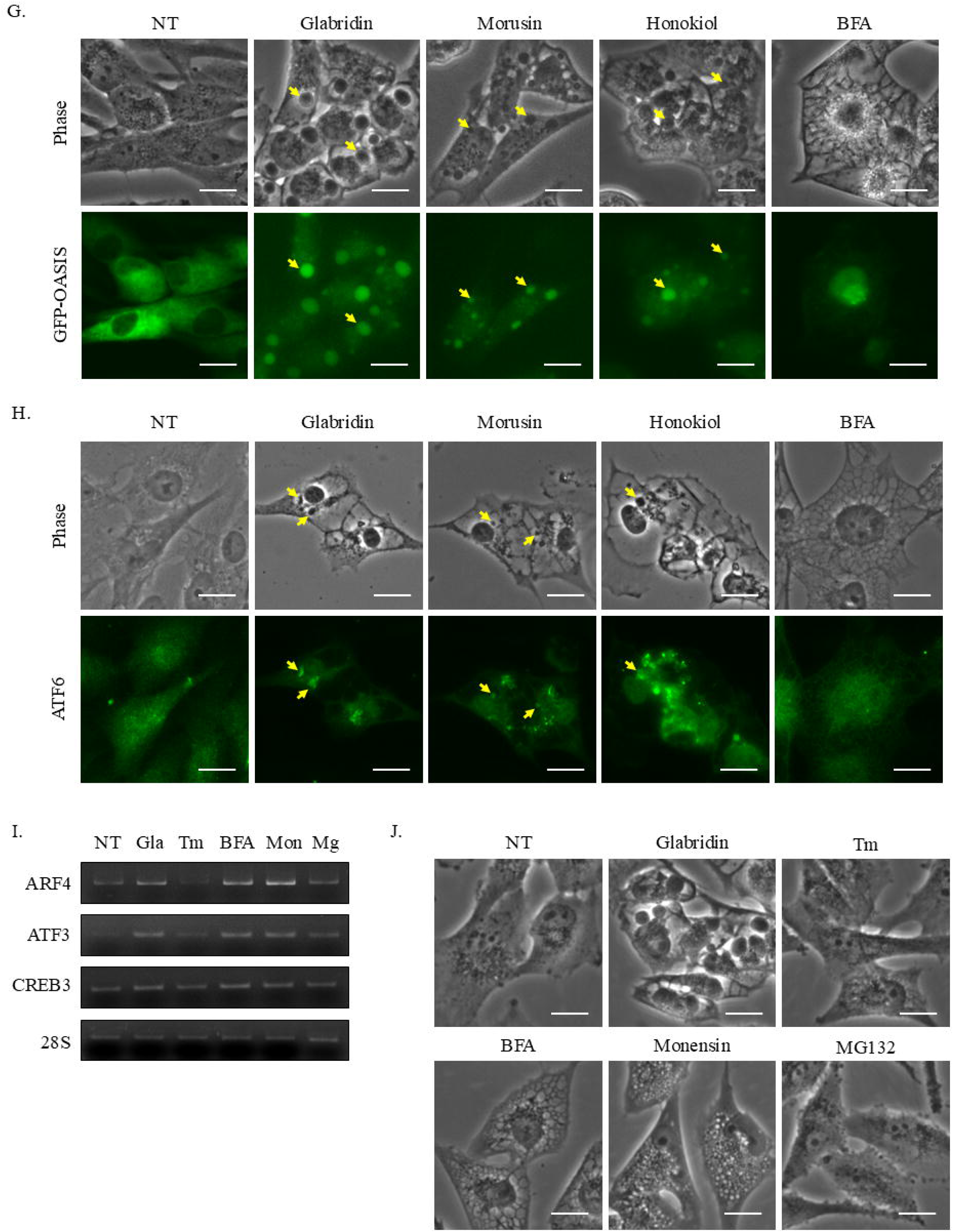
Functional Changes in the Golgi Apparatus and Protein Trafficking During Paraptosis. (A-B) Immunostaining with the cis-Golgi marker GM130 (A) and the cis-/medial Golgi marker Giantin (B) in U251MG cells treated with glabridin (60 μM), morusin (40 μM), honokiol (60 μM), or curcumin (40 μM) for 24 hours, showing disintegration of the typical Golgi structure adjacent to the nucleus. Scale bar: 20 μm. (C) Western blot analysis of GM130 and Syntaxin 16, showing no significant changes in expression levels in cells treated with paraptosis inducers. (D) ELISA analysis indicating a significant reduction in the extracellular secretion of VEGFA protein after treatment with paraptosis inducers. BFA was used as a positive control. Mean ± SD, n=3. *P<0.05 vs. non-treatment group. (E) RT-PCR analysis of Golgi stress markers ARF4, ATF3, and CREB3, showing significant increase in cells treated with glabridin, morusin, and honokiol, but less pronounced effects with curcumin. 28S was used as a loading control. (F) Western blot analysis of OASIS and ATF6 in U251MG cells stably expressing Flag-OASIS. The presence of lower molecular weight bands indicates cleavage of these proteins by paraptosis inducers. F: full length, N: N-terminus. (G) Fluorescence microscopy showing the localization of OASIS in U251MG cells stably expressing GFP-OASIS treated with glabridin (60 μM), morusin (40 μM), honokiol (60 μM), and BFA (1 μM) for 24 hours. Cleaved N-terminal fragments of OASIS accumulate around the nucleus and overlap with putative paraptosomes (yellow arrows). BFA was used as a positive control. Scale bar: 20 μm. (H) Immunostaining of ATF6 in U251MG cells treated with paraptosis inducers, showing bright punctate structures around the nucleus (yellow arrows). (I) RT-PCR analysis of Golgi stress markers ARF4, ATF3, and CREB3 in U251MG cells treated with various stress inducers, including tunicamycin, BFA, monensin, and MG132, indicating differential activation of Golgi stress responses. 28S was used as a loading control. (J) Light microscopy images showing cytoplasmic vacuolation in cells treated with BFA and monensin, without the formation of putative paraptosomes. Tunicamycin and MG132 treatments did not result in notable morphological changes. Scale bars: 20 μm. Tm: Tunicamycin, BFA: Brefeldin A, Mg: Mg132

The endoplasmic reticulum (ER) and Golgi apparatus, which are closely linked and critical for protein synthesis and processing, both exhibited changes during paraptosis. To observe protein transport and processing between these organelles, we utilized the ER stress sensor proteins OASIS and ATF6 [31–32]. These transmembrane proteins typically translocate from the ER to the Golgi apparatus under ER stress, where they are cleaved by Site-1 Protease (S1P) and Site-2 Protease (S2P), releasing an N-terminal fragment that translocates to the nucleus [31–32]. Western blot analysis revealed that both OASIS and ATF6 were cleaved by paraptosis inducers, as evidenced by the appearance of lower molecular weight bands (Figure 5F). Interestingly, fluorescence microscopy showed that the cleaved N-terminal fragment of OASIS did not fully enter the nucleus during paraptosis, but instead accumulated around the nucleus in a circular pattern, overlapping with the putative paraptosomes (Figure 5G). This phenomenon contrasts with cells treated with Brefeldin A (BFA), a known Golgi disassembly agent and ER stress inducer [30], where the N-terminal fragment of OASIS entered the nucleus as expected. Immunostaining of ATF6 revealed a similar pattern, with signals highly concentrated in perinuclear punctate structures, despite the disappearance of putative paraptosomes after paraformaldehyde (PFA) fixation (Figure 5H). These observations indicate significant alterations in the typical trafficking and processing pathways of ER stress-related proteins during paraptosis, pointing to a unique and potentially disruptive mechanism affecting cellular homeostasis.

Given the significant alterations in Golgi morphology and function during paraptosis, we aimed to determine whether Golgi stress alone could induce paraptosis or putative paraptosome formation. Additionally, we investigated the potential contributions of ER stress and proteasome inhibition, which are known to be associated with paraptosis. To this end, we employed specific stress inducers, including tunicamycin for ER stress, BFA and monensin for Golgi stress, and the proteasome inhibitor MG132. RT-PCR analysis showed that BFA and monensin significantly increased the expression of Golgi stress markers ARF4, ATF3, and CREB3, while tunicamycin and MG132 had minimal effects (Figure 5I). Light microscopy revealed that BFA and monensin induced significant cytoplasmic vacuolation with varied morphologies, but notably, neither treatment led to the formation of putative paraptosomes (Figure 5J). Tunicamycin and MG132 treatments did not result in notable morphological changes. Collectively, these results suggest that while ER stress, Golgi stress, and proteasome inhibition are associated with paraptosis, the isolated induction of these stress responses is insufficient to trigger putative paraptosome formation.

## Discussion

Our study unveils a novel aspect of paraptosis: the formation of distinctive high-density spherical structure, which we have termed “paraptosome.” The structure, characterized by a dense network of intertwined membranes, represent a complex reorganization of cellular components. Unlike the low-density vacuoles traditionally associated with paraptosis, these high-density structures emerge as a unique morphological hallmark of the cell death process induced by specific compounds such as glabridin, morusin, and honokiol. The detailed ultrastructure of putative paraptosomes, revealed through electron microscopy, distinguishes them from typical cellular vesicles or organelles. This distinctive morphology suggests that putative paraptosomes could serve as a novel biomarker for paraptosis, potentially enhancing our ability to identify and study this cell death pathway in various cellular contexts.

A key finding of our study is the close association between putative paraptosome formation and Golgi apparatus dysfunction. The transformation of normal Golgi morphology into high-density circular structures, coupled with upregulation of Golgi stress markers ARF4, ATF3, and CREB3, and reduced VEGFA secretion, indicates a profound disruption of Golgi function during paraptosis. The increase in the cleavage of OASIS and ATF6, and the retention of their cleaved N-terminal fragments within these circular structures, rather than their typical nuclear translocation, reveals a novel disruption of ER-Golgi-nucleus signal transduction in paraptosis. Additionally, alterations in LAMP1 protein bands may be attributed to Golgi dysfunction affecting LAMP1 protein modification [33]. These observations extend our understanding of paraptosis beyond the well-established roles of ER stress and proteasome inhibition, highlighting Golgi dysfunction as a potentially central feature of this mode of cell death.

Our study underscores the complex interplay between various organelles and cellular processes in the execution of paraptosis, suggesting a phenomenon akin to multi-organ dysfunction at the cellular level. We observed significant alterations in mitochondrial function, including membrane potential loss, increased ROS production, and decreased Na^+^/K^+^ and Ca^2+^/Mg^2+^ ATPase activities, likely contributing to the energy depletion and ion homeostasis imbalance characteristic of this cell death mode. Additionally, the disruption of ER function and autophagic processes, as evidenced by changes in the expression and fluorescence patterns of LC3 and p62, indicates extensive organelle dysfunction. These multi-faceted cellular stresses may overwhelm the cell’s adaptive responses, ultimately leading to death. Importantly, the inability of isolated ER stress, Golgi stress, proteasome inhibition, or autophagy alterations to trigger putative paraptosome formation supports the notion that paraptosis results from a complex, coordinated stress response involving multiple organelles and signaling pathways. This comprehensive disruption across multiple cellular systems highlights the intricate nature of paraptosis and its distinction from other forms of cell death. This “cellular multi-organ dysfunction” may explain its effectiveness against drug-resistant cancer cells that have developed mechanisms to evade traditional apoptotic pathways.

Curcumin appears to exhibit some distinct effects in our study compared to glabridin, morusin, and honokiol. While curcumin has been reported to induce paraptosis-like cell death in some cell types [8, 11], our observations reveal nuanced differences in its effects in U251MG cells. The curcumin-induced vacuoles showed distinct properties, and the upregulation of Golgi stress markers differed from that observed with other compounds in our study. Additionally, we noted differences in ROS production, LC3B accumulation, and PARP cleavage patterns compared to glabridin, morusin, and honokiol treatments. These observations suggest that the mechanisms of curcumin-induced cell death in U251MG cells may involve additional or alternative pathways alongside paraptosis. This underscores the intricate nature of paraptosis and highlights the importance of considering both cell type-specific and compound-specific effects in paraptosis research. Further investigation into these responses could provide valuable insights into the diverse mechanisms through which different compounds induce paraptosis across various cellular contexts.

Despite these significant findings, several questions remain unanswered, pointing to future research directions. The specific molecular mechanisms of putative paraptosome formation and its role in the paraptosis process require further investigation. One possibility is that paraptosis-induced proteasome inhibition and autolysosome formation blockade lead to the accumulation of proteins in putative paraptosomes, as many key proteins in signaling pathways seem to increase non-specifically during paraptosis. This hypothesis requires further experimental validation. Additionally, our study highlights a significant technical challenge: the sensitivity of putative paraptosomes to paraformaldehyde fixation. This sensitivity complicates immunofluorescence studies and identifying paraptosis in human specimens. Developing alternative fixation methods that preserve putative paraptosome structures is crucial for advancing paraptosis research in clinical contexts. Addressing the disappearance of putative paraptosomes post-fixation is critical for clinical applications, as morphological observations in human specimens typically require fixation. Innovative fixation methods will enable more accurate identification and study of paraptosis in clinical samples, providing insights into its role in human diseases and potential therapeutic interventions. Future research should prioritize developing such methods to bridge laboratory findings and clinical applications.

In conclusion, our study provides new insights into the cellular mechanisms of paraptosis, highlighting the formation of putative paraptosomes as a novel and potentially defining feature of this mode of cell death. The central role of Golgi apparatus dysfunction and the complex interplay between various organelles in this process open up new avenues for understanding and potentially targeting paraptosis. These findings not only advance our fundamental understanding of cell death mechanisms but also may have significant implications for the development of new therapeutic strategies against drug-resistant cancers. Future research focusing on the regulation of putative paraptosome formation, the development of improved detection methods, and the exploration of paraptosis in diverse cellular contexts will be crucial for translating these findings into clinical applications.

## Materials and methods

### Cell culture and reagents

U251MG cells were cultured in Dulbecco’s Modified Eagle’s Medium (11965118, Gibco) supplemented with 10% fetal bovine serum (E600001, Sangon Biotech). Cells were maintained at 37°C in a humidified atmosphere with 5% CO2. Glabridin (S31750) was purchased from Yuanye Bio-Technology. Morusin (GN10436), honokiol (GN10664), curcumin (GC14787), and MG132 (GC10383) were obtained from GLPBIO. Tunicamycin (ab120296) was sourced from Abcam, while brefeldin A (S1536) and monensin (S1753) were acquired from Beyotime Biotechnology.

### Stable cell lines

Plasmids for the expression of fluorescently tagged proteins were constructed using the pCDH-CMV-MCS-EF1-Puro vector system. Constructs were generated to label specific cellular organelles and structures by fusing target sequences to fluorescent proteins. These included KDEL-GFP for the endoplasmic reticulum, COX8-RFP for mitochondria, RFP-LC3B for autophagosomes, LAMP1-RFP for lysosomes, and B4GALT1(1-82)-RFP for the Golgi apparatus. For the expression of GFP-OASIS and Flag-OASIS, plasmids were constructed using the pCDH-CMV-MCS-EF1-Hygro viral vector system. Each construct was inserted into the appropriate vector, confirmed by sequencing, and subsequently packaged into lentiviral particles. U251MG cells were transduced with these lentiviral particles to generate stable cell lines expressing the respective tagged proteins.

### Cell viability assay

The cell counting kit-8 (K1018, APExBIO) assay was utilized to evaluate the cytotoxicity of drugs. Briefly, cells were plated at a density of 5 × 10^3^ cells/well in 96-well plates. They were then exposed to concentrations of 20, 40, 60, 80, and 100 μM of the following compounds: glabridin, morusin, honokiol, and curcumin for a duration of 24 hours. Following the exposure period, 10 μl of CCK-8 reagent was added directly to each well. Due to the dark brown color of curcumin, which could interfere with the optical density readings, the medium was replaced before adding the CCK-8 reagent for the curcumin-treated cells. The plates were subsequently incubated for an additional 2 hours at 37°C. After incubation, the optical density (OD) at 450 nm was measured using a microplate reader (Infinite M Nano, TECAN) to determine the cytotoxic effects of the compounds. respective tagged proteins.

### Flow cytometry

Cells were seeded in a 10 cm dish at a density of 1× 10^6^ cells and treated with 60 μM glabridin, 40 μM morusin, 60 μM honokiol, or 40 μM curcumin for 24 hours. The cells were then harvested, washed with 1× PBS, and fixed in 70% ethanol at -20°C for 2 hours. After fixation, the cells were centrifuged and washed with 1× PBS twice. One unit of RNase A was added to the cell suspension and incubated for 30 minutes at 37°C. Subsequently, the cells were resuspended in 300 μl of PI staining buffer and incubated at 37°C for 10 minutes before analysis. The DNA content of the cells was then analyzed by flow cytometry (Novocyte Quanteon, Agilent).

### RT-PCR

Total RNA was extracted from cells using TRNzol reagent (DP405-02, TIANG EN) following the manufacturer’s instructions. First-strand cDNA was synthesiz ed in a 20 μl reaction volume using random primers (3802, TAKARA) and 1 μl M-MLV-RT enzyme (M1701, Promega). PCR was performed in a 20 μl vol ume containing 10 pmol of each primer, 4 μmol dNTPs (PC2300, Solarbio), 1 unit of Taq DNA polymerase (4992763, TIANGEN), and 1× PCR buffer. The primer sequences are as follows: 28S Fwd 5’-TTGAAAATCCGGGGGAGAG-3’, Rev 5’-ACATTGTTCCAACATGCCAG-3’; BiP Fwd 5’-GTTTGCTGAGGA AGACAAAAAGCTC-3’, Rev 5’-CACTTCCATAGAGTTTGCTGATAAT-3’; XB P1 Fwd 5’-GTTGAGAACCAGGAGTTAAG-3’, Rev 5’-GGTGACAACTGGGCC TGCAC-3’; CHOP Fwd 5’-GGAAACAGAGTGGTCATTCCC-3’, Rev 5’-CTGC TTGAGCCGTTCATTCTC-3’; ARF4 Fwd 5’-CCCTCTTCTCCCGACTATTTGG-3’, Rev 5’-GCACAAGTGGCTTGAACATACC-3’; ATF3 Fwd 5’-GCTGTCACC ACGTGCAGTAT-3’, Rev 5’-TTTGTGTTAACGCTGGGAGA-3’; CREB3 Fwd 5’ -AAAGTGGAGATTTGGGGACG-3’, Rev 5’-CGCTCGGTACCTCAGAAAG-3’. PCR products were analyzed by electrophoresis on a 4.8% polyacrylamide gel and visualized using a gel documentation system. The expression levels of targ et genes were normalized to the expression of the 28S rRNA gene as an inter nal control.

### Western Blot Analysis

Cells were harvested and lysed in RIPA buffer (P0013B, Beyotime) supplemented with Protease Inhibitor Cocktail (K1010, APExBIO). Protein concentrations were determined using the BCA Protein Assay Kit (P0012, Beyotime). Equal amounts of protein samples (10-20 μg) were separated by SDS-PAGE and transferred onto PVDF membranes (IPVH00010, Millipore). Membranes were blocked with 5% non-fat milk in TBS-T (Tris-buffered saline with 0.1% Tween-20) for 1 hour at room temperature and then incubated overnight at 4°C with primary antibodies against GAPDH (T0004, Affinity), cleaved caspase-3 (AF7022, Affinity), cleaved PARP (ASP214, Affinity), ubiquitin (H1021, Santa Cruz), LAMP1 (A16894, ABclonal), LC3 A/B (AF5402, Affinity Biosciences), p62/SQSTM1 (bs-55207R, Bioss), GM130 (A5344, ABclonal), Syntaxin 16 (DF12330, Affinity), Flag (AF2852, Beyotime), ATF6 (A0202, ABclonal), MFF (A4874, ABclonal), and phospho-DRP1 (Ser637) (DF2980, Affinity). After washing with TBS-T, membranes were incubated with HRP-conjugated secondary antibodies (bs-0294R-HRP and bs-0296G-HRP, Bioss) for 1 hour at room temperature. Protein bands were detected using an ECL detection system (ABL-X5, Tanon).

### Mitochondrial membrane potential assay

Mitochondrial membrane potential was measured using JC-1 staining (C2005, Beyotime). After treating cells with 60 μM glabridin, 40 μM morusin, 60 μM honokiol, or 40 μM curcumin for 6 h, the cells were incubated with 10 μg/ml JC-1 for 30 minutes at 37°C. The stained cells were then visualized using a fluorescence microscope (EVOS M5000, Invitrogen).

### Reactive oxygen species assay

Cells were grown on coverslips in 24-well plates for 24 hours and then treated with 60 μM glabridin, 40 μM morusin, 60 μM honokiol, or 40 μM curcumin for 6 hours. After treatment, the cells were incubated with 10 μM DCFH-DA (BL714A, Biosharp) for 30 minutes. Following incubation, the cells were imaged using a fluorescence microscope (EVOS M5000, Invitrogen).

### ATPase assay

Na^+^/K^+^ ATPase and Ca^2+^/Mg^2+^ ATPase activities were measured using specific ATPase Assay Kits (BC0065 and BC0965, Solarbio) according to the manufacturer’s instructions. U251MG cells were treated with 60 μM glabridin, 40 μM morusin, 60 μM honokiol, or 40 μM curcumin for 24 hours. After treatment, cells were harvested and homogenized in the provided assay buffer. The homogenates were centrifuged at 12,000 g for 15 minutes at 4°C, and the supernatants were collected for ATPase activity measurement.

### Time-laps imaging

Time-lapse imaging was performed to monitor the dynamic changes in Golgi structures during paraptosis. U251MG cells stably expressing the Golgi-specific fluorescent marker B4GALT1(1-82)-RFP were seeded in a glass-bottom dish (BS-20-GJM, Biosharp) and treated with 60 μM glabridin. Imaging was initiated immediately after drug treatment and continued for 5 hours. Images were captured every 3 minutes using a confocal laser scanning microscope (Nikon AX) equipped with a 60× oil immersion objective.

### Transmission electron microscopy

Cells were treated with 60 μM glabridin for 24 h, and initially fixed with 2.5% glutaraldehyde for 1 h. Then the cells were postfixed in 1% osmium tetroxide for 1 hat 4 °C. After dehydration with graded ethanol series, the cells were embedded in EMBED 812, polymerized, and observed under the electron microscope (HT7800, HITACHI).

### Elisa assay

U251MG cells were treated with 60 μM glabridin, 40 μM morusin, 60 μM honokiol, 40 μM curcumin, or 1 μM BFA for 24 hours. After treatment, culture media were collected, centrifuged to remove debris, and analyzed for VEGFA concentration using the ELISA kit (RX105007H, RUIXIN Biotech) according to the manufacturer’s instructions. The optical density (OD) was measured at 450 nm using a microplate reader (Infinite M Nano, TECAN).

### Statistical analyses

Data are presented as mean ± SD from at least three independent experiments for each experimental condition. Statistical significance was determined using Student’s t-test to calculate P values, with P < 0.05 considered statistically significant.

## Declarations

### Competing interests

The authors declare no competing interests.

### Author contributions

Xiang Cui and Min Cui co-designed and performed the experiments. Xiang Cui organized the data and wrote the manuscript. Min Cui provided essential guidance and contributed to the data analysis and interpretation. Hongda Zheng, Fang Zhang, and Liao Yang assisted with the experimental work and data analysis. Haoming Li provided technical support and contributed to the interpretation of the data. Jiayu Ni, Dengfeng Wang, Huali Zhang, Pan Tang, and Ru Li contributed to the preparation of reagents and provided experimental support. Qi Zhang provided critical feedback and offered valuable insights into the interpretation of the results.

## Acknowledgments

This research was supported by Guangxi Natural Science Foundation of China (2021GXNSFBA220005), the Project for Enhancing Young and Middle-aged Teacher’s Research Basis Ability in Colleges of Guangxi (No. 2024KY0517), and the “13th Five-Year” Science and Technology Research Project of Jilin Provincial Education Department (JJKH20191142KJ). Min Cui, Hongda Zheng, and Qi Zhang were supported by the Guangxi Medical and Health Key Cultivation Discipline Construction Project.

Declaration of generative AI and AI-assisted technologies in the writing process

During the preparation of this work the authors used ChatGPT to improve language and readability. After using this tool, the authors reviewed and edited the content as needed and takes full responsibility for the content of the publication.

## Notes

### Competing Interest Statement

The authors have declared no competing interest.

## References

[1] Park W, Wei S, Kim BS, et al. (2023). Diversity and complexity of cell death: a historical review. Exp Mol Med, 55(8), 1573–1594.

[2] Hanson S, Dharan APVJ, et al. (2023). Paraptosis: a unique cell death mode for targeting cancer. Front Pharmacol, 14, 1159409.

[3] Sperandio S, de Belle I, Bredesen DE. (2000). An alternative nonapoptotic form of programmed cell death. Proc Natl Acad Sci U S A, 97(26), 14376–81.

[4] Kim E, Lee DM, Seo MJ, Lee HJ, Choi KS. (2021). Intracellular Ca2+ imbalan ce critically contributes to paraptosis. Front Cell Dev Biol, 8, 607844.

[5] Cui X, Cui M. (2022). Glabridin induces paraptosis-like cell death via ER stress in breast cancer cells. Heliyon, 8(9), e10607.

[6] Xue J, Li R, Zhao X, et al. (2018). Morusin induces paraptosis-like cell death t hrough mitochondrial calcium overload and dysfunction in epithelial ovarian canc er. Chem Biol Interact, 283, 59–74.

[7] Liu X, Gu Y, Bian Y, et al. (2021). Honokiol induces paraptosis-like cell death of acute promyelocytic leukemia via mTOR & MAPK signaling pathways activat ion. Apoptosis, 26(3-4), 195–208.

[8] Garrido-Armas M, Corona JC, Escobar ML, et al. (2018). Paraptosis in human g lioblastoma cell line induced by curcumin. Toxicol In Vitro, 51, 63–73.

[9] Sperandio S, Poksay K, de Belle I, et al. (2004). Paraptosis: mediation by MAP kinases and inhibition by AIP-1/Alix. Cell Death Differ, 11(10), 1066–75.

[10] Liu S, Tian Y, Liu C, Gui Z, Yu T, Zhang L. (2024). TNFRSF19 promotes end oplasmic reticulum stress-induced paraptosis via the activation of the MAPK path way in triple-negative breast cancer cells. Cancer Gene Ther, 31(2), 217–227.

[11] Yoon MJ, Kim EH, Lim JH, Kwon TK, Choi KS. (2010). Superoxide anion and proteasomal dysfunction contribute to curcumin-induced paraptosis of malignant breast cancer cells. Free Radic Biol Med, 48(5), 713–26.

[12] Ma L, Xuan X, Fan M, et al. (2022). A novel 8-hydroxyquinoline derivative ind uces breast cancer cell death through paraptosis and apoptosis. Apoptosis, 27(7-8), 577–589.

[13] Yoon MJ, Lee AR, Jeong SA, et al. (2014). Release of Ca2+ from the endoplas mic reticulum and its subsequent influx into mitochondria trigger celastrol-induce d paraptosis in cancer cells. Oncotarget, 5(16), 6816–31.

[14] Dilshara MG, Neelaka Molagoda IM, Prasad Tharanga Jayasooriya RG, Choi YH, Park C, Kim GY. (2021). Indirubin-3’-monoxime induces paraptosis in MDA-M B-231 breast cancer cells by transmitting Ca2+ from endoplasmic reticulum to m itochondria. Arch Biochem Biophys, 698, 108723.

[15] Bury M, Girault A, Mégalizzi V, et al. (2013). Ophiobolin A induces paraptosis-like cell death in human glioblastoma cells by decreasing BKCa channel activity. Cell Death Dis, 4(3), e561.

[16] Yumnam S, Hong GE, Raha S, et al. (2016). Mitochondrial Dysfunction and Ca (2+) Overload Contributes to Hesperidin Induced Paraptosis in Hepatoblastoma C ells HepG2. J Cell Physiol, 231(6), 1261–8.

[17] Nedungadi D, Binoy A, Vinod V, et al. (2021). Ginger extract activates caspase independent paraptosis in cancer cells via ER stress, mitochondrial dysfunction, AIF translocation, and DNA damage. Nutr Cancer, 73(1), 147–159.

[18] Binoy A, Nedungadi D, Katiyar N, et al. (2019). Plumbagin induces paraptosis i n cancer cells by disrupting the sulfhydryl homeostasis and proteasomal function. Chem Biol Interact, 310, 108733.

[19] Hager S, Korbula K, Bielec B, et al. (2018). The thiosemicarbazone Me2NNMe2 induces paraptosis by disrupting the ER thiol redox homeostasis based on prote in disulfide isomerase inhibition. Cell Death Dis, 9(11), 1052.

[20] Seo MJ, Lee DM, Kim IY, et al. (2019). Gambogic acid triggers vacuolation-ass ociated cell death in cancer cells via disruption of thiol proteostasis. Cell Death Dis, 10(3), 187.

[21] Chen X, Chen X, Zhang X, et al. (2019). Curcuminoid B63 induces ROS-media ted paraptosis-like cell death by targeting TrxR1 in gastric cells. Redox Biol, 21, 101061.

[22] Chen Q, Song S, Wang Z, et al. (2021). Isorhamnetin induces the paraptotic cell death through ROS and the ERK/MAPK pathway in OSCC cells. Oral Dis, 27 (2), 240–250.

[23] Nguyen PL, Lee CH, Lee H, Cho J. (2022). Induction of Paraptotic Cell Death in Breast Cancer Cells by a Novel Pyrazolo[3,4-h]quinoline Derivative through R OS Production and Endoplasmic Reticulum Stress. Antioxidants (Basel), 11(1), 1 17.

[24] Zhang JS, Li DM, He N, et al. (2011). A paraptosis-like cell death induced by δ-tocotrienol in human colon carcinoma SW620 cells is associated with the supp ression of the Wnt signaling pathway. Toxicology, 285(1-2), 8–17.

[25] Suresh RN, Jung YY, Harsha KB, Mohan CD, Ahn KS, Rangappa KS. (2024). Isoxazolyl-urea derivative evokes apoptosis and paraptosis by abrogating the Wnt/β-catenin axis in colon cancer cells. Chem Biol Interact, 399, 111143.

[26] Almanza A, Carlesso A, Chintha C, et al. (2019). Endoplasmic reticulum stress signalling - from basic mechanisms to clinical applications. FEBS J, 286(2), 241–278.

[27] Khoder-Agha F, Harrus D, Brysbaert G, et al. (2019). Assembly of B4GALT1/S T6GAL1 heteromers in the Golgi membranes involves lateral interactions via hig hly charged surface domains. J Biol Chem, 294(39), 14383–14393.

[28] Manickam V, Tiwari A, Jung JJ, et al. (2011). Regulation of vascular endothelial growth factor receptor 2 trafficking and angiogenesis by Golgi localized t-SNA RE syntaxin 6. Blood, 117(4), 1425–35.

[29] Reiling JH, Olive AJ, Sanyal S, et al. (2013). A CREB3-ARF4 signalling pathw ay mediates the response to Golgi stress and susceptibility to pathogens. Nat Cel l Biol, 15(12), 1473–85.

[30] Oh-Hashi K, Hasegawa T, Mizutani Y, Takahashi K, Hirata Y. (2021). Elucidati on of brefeldin A-induced ER and Golgi stress responses in Neuro2a cells. Mol Cell Biochem, 476(10), 3869–3877.

[31] Murakami T, Kondo S, Ogata M, et al. (2006). Cleavage of the membrane-boun d transcription factor OASIS in response to endoplasmic reticulum stress. J Neur ochem, 96(4), 1090–100.

[32] Ye J, Rawson RB, Komuro R, et al. (2000). ER stress induces cleavage of me mbrane-bound ATF6 by the same proteases that process SREBPs. Mol Cell, 6(6), 1355–64.

[33] Baba K, Kuwada S, Nakao A, et al. (2020). Different localization of lysosomal-associated membrane protein 1 (LAMP1) in mammalian cultured cell lines. Histo chem Cell Biol, 153(4), 199–213.

