## Supplementary figures and images for "Paraptosome: A Novel Pathological Feature in Paraptotic Cell Death"

### Supplementary Figure1

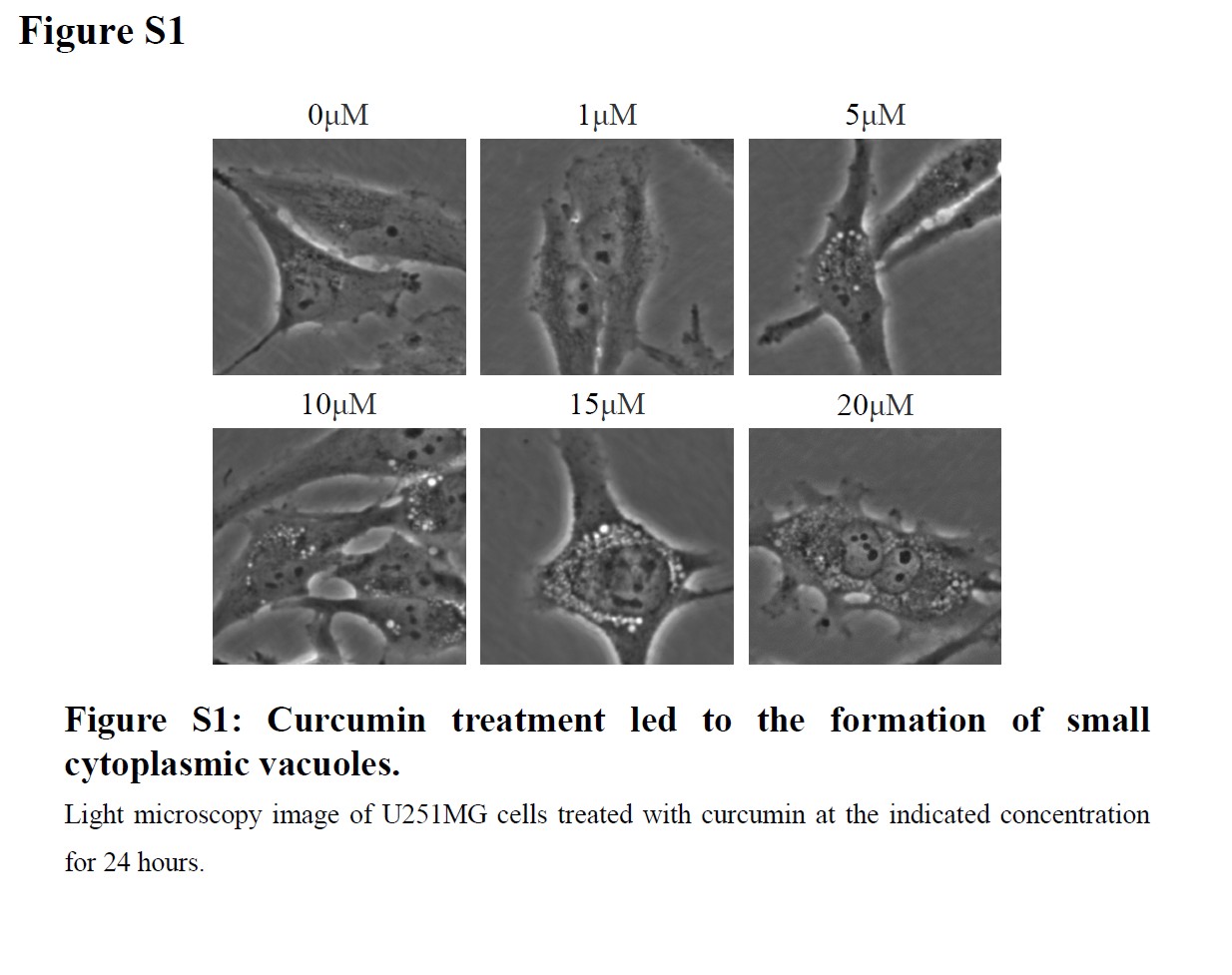
